# Ecosystem Entanglement and the Propagation of Nutrient-Driven Instability

**DOI:** 10.1101/2020.04.20.050302

**Authors:** Kevin S. McCann, A.S MacDougall, G.F. Fussmann, C. Bieg, K. Cazelles, M.E. Cristescu, J.M. Fryxell, G. Gellner, B. Lapointe, A. Gonzalez

## Abstract

Almost 50 years ago, Michael Rosenzweig pointed out that nutrient addition can destabilize food webs, leading to loss of species and reduced ecosystem function through the paradox of enrichment. Around the same time, David Tilman demonstrated that increased nutrient loading would also be expected to cause competitive exclusion leading to deleterious changes in food web diversity. While both concepts have greatly illuminated general diversity-stability theory, we currently lack a coherent framework to predict how nutrients influence food web stability across a landscape. This is a vitally important gap in our understanding, given mounting evidence of serious ecological disruption arising from anthropogenic displacement of resources and organisms. Here, we combine contemporary theory on food webs and meta-ecosystems to show that nutrient additions are indeed expected to drive loss in stability and function in human-impacted regions. However, this loss in stability occurs not just from wild oscillations in population abundance, but more frequently from the complete loss of an equilibrium due to edible plant species being competitively excluded. In highly modified landscapes, spatial nutrient transport theory suggests that such instabilities can be amplified over vast distances from the sites of nutrient addition. Consistent with this theoretical synthesis, the empirical frequency of these distant propagating ecosystem imbalances appears to be growing. This synthesis of theory and empirical data suggests that human modification of the Earth’s ecological connectivity is “entangling” once distantly separated ecosystems, causing rapid, expansive, and costly nutrient-driven instabilities over vast areas of the planet. The corollary to this spatial nutrient theory, though -- akin to weak interaction theory from food web networks -- is that slow spatial nutrient pathways can be potent stabilizers by moderating flows across a landscape.

## Main Text

The growing demand for food is producing ecologically homogenized agroecosystems that now dominate over one third of the Earth’s habitable land surface (Bruinsma & FAO 2003; Nyström *et al*. 2019). Human wastewater, fossil fuel emissions, deforestation and agricultural intensification for crops and livestock have vastly increased nutrient flows between terrestrial and aquatic ecosystems (Fig. 1A-D). Because homogenized agricultural landscapes lack the extended storage capacity formerly provided by extensive networks of wetlands, nutrients flow rapidly from crop fields, often equipped with tile drainage, to channelized streams before reaching rivers, lakes and oceans. Collectively, these landscape modifications increase and accelerate the movement of nutrients across the land and water (Bennett *et al*. n.d.; Raymond *et al*. 2008; Aufdenkampe *et al*. 2011; Elser & Bennett 2011; Nyström *et al*. 2019). This heightened connectivity means that the impacts of local activity rapidly propagate ‘downstream’, causing farm nutrients and chemicals to accumulate in vast quantities in lakes and oceans where other movement vectors such as ocean currents distribute them globally.

**Fig. 1.**
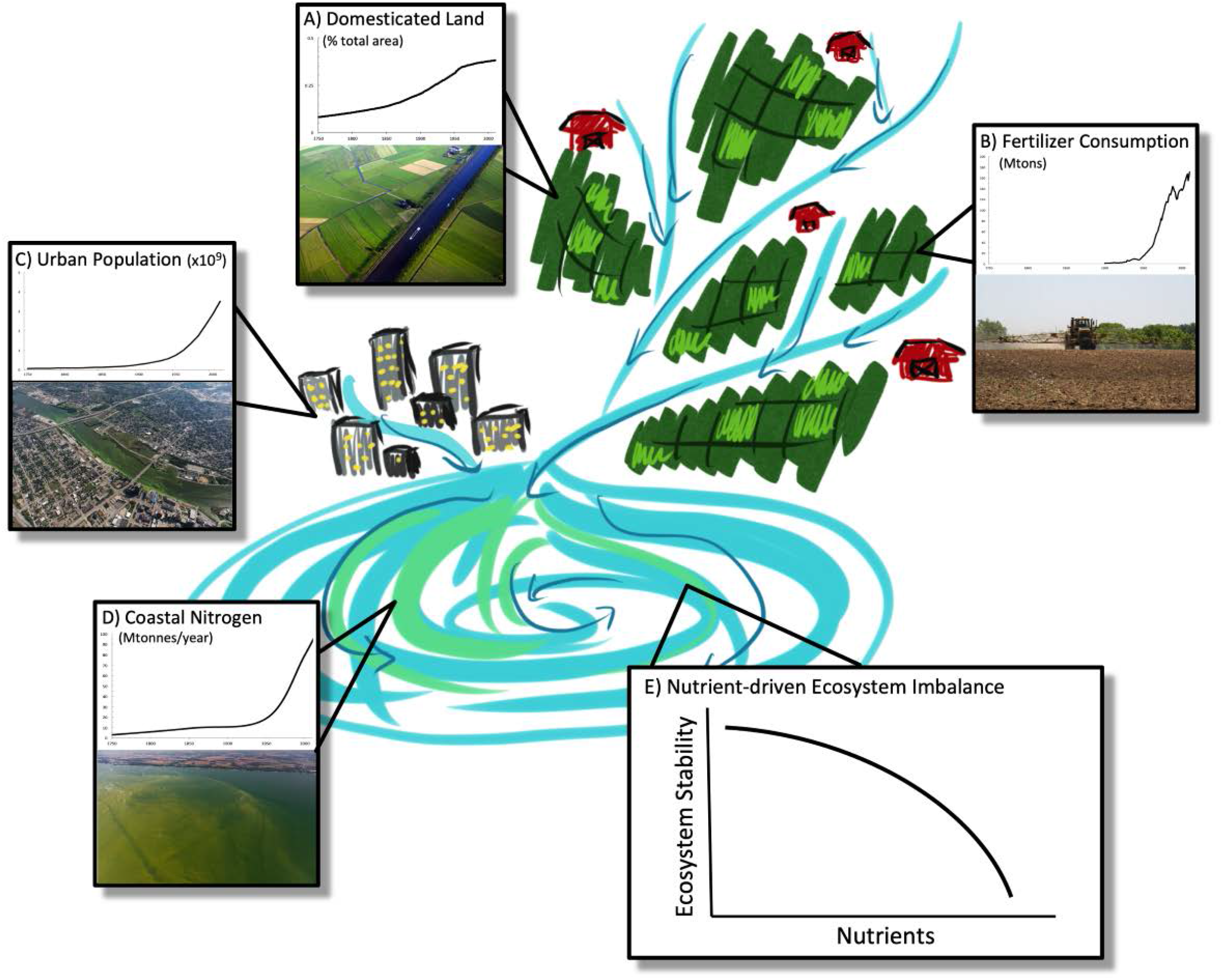
Picture of an increasingly common set of multiscale influences on ecosystems. Natural land is being developed into agricultural land at an increasing rate (inset A). A highly agricultural landscape that removes natural spatial heterogeneity as it is made into one large agro-ecosystem producing a homogenized regional scale system. Globally this landscape modification has been accompanied by an increasing use of nutrients on farms (inset B). Increasing human population is increasing urban populations globally (inset C), and land modification on generic farms reduce or remove riparian buffers, tile drains fields, and channelizes streams (inset A) -- all activities that ramp up the loading and movement of sediment including nutrients from the field to the creeks, to the streams, rivers, and finally lakes and oceans where nutrients accumulate (inset D). This threatens the ecological balance of these terminal ecosystems (i.e., instability takes place at a distance from local actions), often with nutrient-driven instabilities such as harmful algal blooms and dead zones (inset E). The form of instability can be variance-driven or mean-driven instability (Box 1 and 2). Data from: Steffen et al. 2015. Great Acceleration Data http://www.igbp.net/globalchange/greatacceleration.4.1b8ae20512db692f2a680001630.html Photo credits: A) Peter Prokosch http://www.grida.no/resources/1698 B) P177 https://www.flickr.com/photos/48722974@N07/4478367887 C) Aerial Associates Photography, Inc. by Zachary Haslick D) Aerial Associates Photography, Inc. by Zachary Haslick

The spatial homogenization of landscapes broadly increases across-ecosystem connections -- a form of environmental globalization that we refer to as “ecosystem entanglement”, whereby distant ecosystems become increasingly connected, such that anthropogenic disturbance in one location can have rapid, and often amplified, impacts elsewhere (see landscape example in Fig. 1; Box 1 for definitions). These flows, for example, are now understood to be responsible for massive algal blooms resulting in ecosystem dead zones and are striking examples of regional ecological degradation and instability (Diaz & Rosenberg 2008; Huisman *et al*. 2018; Ho *et al*. 2019). Nutrient additions have increased globally, with ~2-fold increases in reactive nitrogen (N) and phosphorus (P) use over the last half century (Galloway *et al*. 2008; Cordell *et al*. 2009; Macdonald *et al*. 2011; Steffen *et al*. 2015) (Fig. 1B) resulting in increasing nitrogen loads to lakes and coastal oceans (Fig. 1D). These trends promise to continue, with an estimated 50% increase in food demands by 2050 as the human population exceeds 9 billion (Vitousek *et al*. 2009; Eisenhauer *et al*. 2012; Springmann *et al*. 2018). As the problem intensifies, so does the need to understand how it might unfold and why the magnitude of impact is so powerful, both of which inform potential avenues for remediation. To advance understanding we link three fundamental bodies of ecological theory: consumer-resource dynamics due to the Paradox of Enrichment (Rosenzweig 1971), species replacement due to nutrient-mediated competitive exclusion (Tilman 1982), and spatial dynamics represented by recent advances in meta-ecosystem theory (Loreau et al. 2003; Gounand et al. 2018).

### Box. 1 Key Definitions

**Telecoupling** – refers to how connections between nature and human beings are growing ever tighter in a more globalized world in ways that positively and negatively impact humanity.

**Ecosystem entanglement** – a specific form of telecoupling, where large-scale spatial connectivity caused by spatial homogenization and increased material or organismal flows results in a dramatically heightened cross-ecosystems connectivity.

**Dendritic river network** – branching pattern of connections that directionally transports water across landscapes

**Instability** – any form of dynamic that threatens the persistence of a species (i.e., a species reaches dangerously low densities), often measured by coefficient of variation (CV = σ/μ) or strength of attraction to an equilibrium in theoretical models. Local minima on an attractor allow one to assess stability as they detail when populations are at dangerously low densities from oscillations as well as follow the loss of an equilibrium. For more detailed defintions and concpets behind a general loss in stability of a dynamical system see Box.2.

**Equilibrium** – a steady state where all densities are balanced such that gains equal losses.

**Attractor** – an equilibrium or non-equilibrium state (e.g., cycle) to which solutions are attracted after a transient.

**Bifurcation** - a **bifurcation** occurs when a small smooth change made to the parameter values (the **bifurcation** parameters) of a system causes a sudden ‘qualitative’ or topological change in its behavior.

**Cyclic or variance-driven instability**– variability driven by population cycles with potential for a species to approach low densities and go locally extinct at the cycle’s minima. Caused generically by **Hopf** bifurcations in dynamical systems models, where a stable equilibrium turns into limit cycles.

**Structural or Mean-driven instability** – decreasing densities of species, or set of species, which results in the eventual total loss of an equilibrium (i.e., a species mean density declines to zero). Even in simple ecosystem models nutrient increases tend to drive a fundamental change in ecosystem structure that re-routes whole carbon pathways (e.g., increased inedible resources, increased detritus). All such changes are driven by losses of key species that tend to be caused by **transcritical, saddle node or pitchfork** bifurcations in dynamical systems models. In ecological models, transcritical bifurcations are generic for loss of a species, although ecosystem feedbacks can also readily yield saddle node bifurcations that lead to alternate states. See Box. 2 for more details.

**Phase transition** – a qualitative change in the state of a system (e.g. equilibrium) under a continuous change in an external parameter (e.g., increased nutrient flux, and/or nutrient loading).

**Food web or ecosystem module** – building blocks of a whole food web or ecosystem model, generally describing the simplified structure of interactions between major components or guilds within the system. Here, the base ecosystem module includes a consumer, a resource, detritus and nutrients and includes nutrient recycling.

Almost 50 years ago, Michael Rosenzweig warned that increased nutrient loading and higher resultant productivity could lead to ecosystem destabilization and species loss (Rosenzweig 1971) – the exact challenges we are seeing today. This concept, known as the paradox of enrichment (hereafter POE), centered on the irony that the initial benefit of nutrients for higher productivity eventually destabilizes an ecosystem by disrupting the interaction between consumers and their resources. In simple systems with a single species of consumer and a single species of prey (resource), high levels of nutrient loading drive extreme cycles in abundance. At high enough enrichment the oscillatory dynamics cause one species or the other to dip to dangerously low population densities and would therefore go locally extinct in a stochastic world. This oscillatory dynamic has been referred to as *oscillatory or variance-driven instability* (Gellner *et al*. 2016; see Box 2 for details). This mechanism has been verified by highly-controlled trials in some simple experimental microcosms (Luckinbill 1973; 1979; Fussmann et al. 2000), but is rarely confirmed by empirical research in more complex natural settings (although see Tilman and Wedin 1991). Instead, empirical results suggest shifts in ecosystem states without oscillations (Morin & Lawler 1995; Isbell *et al*. 2013). Here, the effects of nutrient loading appear to favor dominance by some species (e.g., harmful algal blooms; HABs) that push the mean densities of other organisms to functional extinction. In food webs this process involves competitive displacement of one kind of prey resource by a second variety that is better adapted to high nutrient conditions (Tilman 1982). We refer to this as *structural or mean-driven ecosystem instability* as it tends to restructure whole carbon pathways in ecosystem models, see Box. 2; (Huisman *et al*. 2018; Ho *et al*. 2019). This common field result suggests a second mechanism for ecological destabilization through enrichment, broadening Rosenzweig’s initial concept of the paradox of enrichment.

### Box. 2 Enrichment and Instability

#### Two General forms of Loss of Stability: Variance (Cyclic) and Mean-driven (structural) losses in stability

Broadly speaking, stability can be lost in an ecological model via wild oscillations or the complete loss of an equilibrium (Fig. B.1A). In 1971, Michael Rosenzweig pointed out that numerous models produced something he coined the paradox of enrichment (Rosenzweig 1971). The idea was simple, he found that changes in carrying capacity (a surrogate for increased nutrients), all else being equal, tended to alter the structure of consumer-resource interactions such that they drove wildly oscillating population dynamics (via a **Hopf bifurcation**; Fig.B.2.A). During these wild cycles (say where nutrient levels are at the red star), the argument goes, population densities of either consumer or resource occasionally plunge to near zero density (see dashed red line which is the persistence line Fig. B.2.CB for example), which increases the likelihood of consumer or resource collapse in a stochastic world (e.g., drought). The paradox therefore was seen as increased nutrients, somewhat counterintuitively, could lead to collapse of C or R or both. This loss of stability has been more recently referred to as **variance-driven or cyclic instability** for the simple reason that it would be expected to largely inflate the standard deviation in a coefficient of variation (referred to as CV in literature) stability calculation (where CV = SD/μ).

Clearly this variance-driven route to instability is not the only form of instability that threatens the persistence of interacting species. Generally speaking, as we vary some parameter there is the possibility that instead of oscillations the varying parameter can push an internal equilibrium (i.e., all species > 0 densities; Fig. B.2.B) to a situation where one species is excluded (i.e., the mean density of one species goes to zero as for the time series example where nutrients are at the red star level; Fig. B.2.D). Since the interior equilibrium is lost, system level stability is clearly breached. This tends to occur via several possible bifurcations, most notably **transcritical or saddle node** in ecology (Grover 1995; Scheffer et al. 2001). Note, since in the above example one of the species at equilibrium (i.e., not deterministically varying over time) is pushed to zero densities this is an example of **mean-driven** loss in stability (CV inflates as mean goes to zero) and signals the complete **loss of an interior equilibrium**. Often, as in the edible-inedible resource case discussed below, this can completely alter the structure of energy flow in the ecosystem.

**Fig. B.2.**
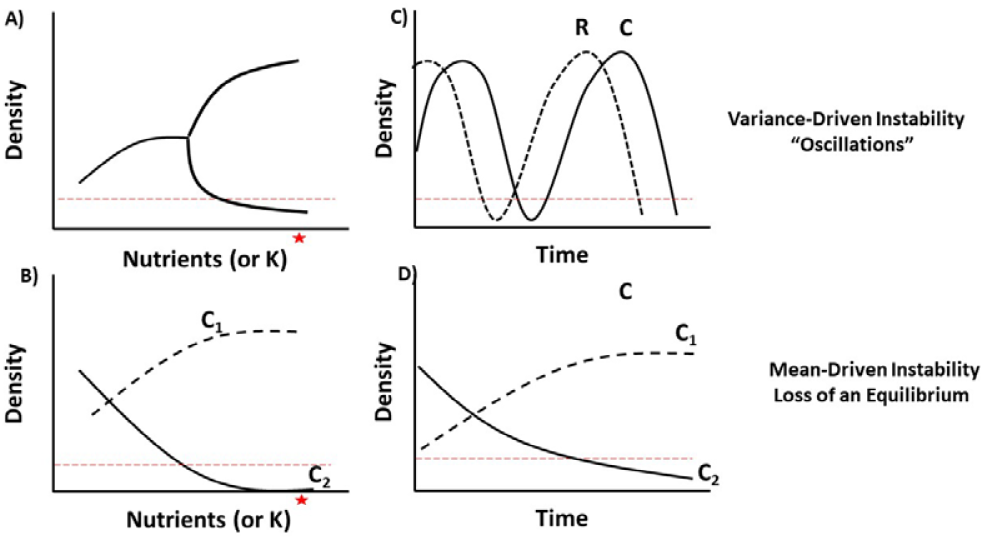
Time series of two different instabilities (loss of persistence of an interaction or set of interactions). A) of an oscillatory loss of stability (variance-driven as mean is high) produced by a deterministic or stochastic Hopf, and; B) a mean-driven loss of stability associated with the complete loss of an equilibrium through a bifurcation such as a transcritical or saddle node.

Although ecology has historically sought to explain ecosystem stability through local processes, recent meta-ecosystem theory suggests that coupling ecosystems across large spatial scales by organismal migration and material transport may play a key role in determining ecosystem stability (Levin 1995; Callaway & Hastings 2002; Gounand *et al*. 2014; Marleau *et al*. 2014; Gravel *et al*. 2016). With the growing empirical evidence of heightened connectivity of nutrients, POE can now be recast as a more general nutrient theory for global ecology. This recast version of POE integrates multiple scales of ecosystem connectivity allowing us to better explain some of the most striking examples of ecosystem imbalance and degradation. One important feature is that increased ecosystem connectivity often occurs in dendritic river networks, where flows are directional, and nutrient concentrations amplify dangerously downstream. The dendritic network is effectively “space filling”, and like a lung, efficiently transports localized inputs (e.g., particle-bound P) into distant waterbodies before ultimately ending in oceans where global current systems move them over vast scales. Our empirical understanding suggests that local land modification enhances the already significant connectivity of directional transport networks (Raymond *et al*. 2008), intensifying ecosystem entanglement and accelerating the build-up of nutrients that destabilize biological processes at great distances (Diaz & Rosenberg 2008).

Here, we spatially extend ecosystem theory to examine how human-mediated increases in nutrient transport can drive widespread long-distance ecosystem instability by integrating POE dynamics and nutrient-mediated competitive exclusion with elevated connectivity (i.e., ‘entanglement’). We define instability as any dynamic outcome that threatens the persistence of species (i.e., via variance or mean-driven ecosystem instability; detailed in Box 2) and alters ecosystem processes due to shifts in the structure and function of food webs. We first argue that a synthesis of existing theory suggests that nutrient enhancement should commonly generate ecosystem instability through either cycle-driven or structural changes in ecological equilibria. To highlight this, we then employ a simple set of ecosystem models with advective **movement of either nutrients or consumers** within a simple dendritic network. We find that increases in nutrient transport (e.g., tile drainage, channelization, climate-derived extreme pulses of local precipitation) or large-scale consumer movement (e.g., ocean circulation, migration) amplify the likelihood of ecological instability (e.g., detrital take over). We then discuss empirical examples that suggest this phenomenon is happening and appears to be increasing globally. We end by suggesting that management of connectivity within directional dendritic networks (Peterson *et al*. 2013a; Galiana *et al*. 2018) could mitigate these costly ecological imbalances at regional and global scales. Specifically, our theory suggests that well-placed landscape scaled management practices (e.g., ‘slow nutrient pathways’ within a spatial network) can act as potent stabilizers of distant ecosystem imbalance, in a manner akin to the stabilizing role weak interactions play in food web theory (May 1973; Gellner and McCann 2016).

## A Brief Review of Nutrient-Stability Theory

Rosenzweig suggested that increasing carrying capacity, a surrogate for nutrient enrichment, can drive population instability by transforming a stable equilibrium into a cyclically unstable one (termed a Hopf bifurcation; see Box 2), such that steady state dynamics give way to consumer-resource oscillations (Rosenzweig 1971). More recent elaborations of food web theory largely agree with this insight (McCann 2011; Murdoch, Briggs, and Nisbet 2013), with one important caveat – any biological structure or process that reduces energy flow between consumer and resource can inhibit the expression of the cyclic instability (e.g., Rip & McCann 2011). As an example, any form of interference that reduces consumer growth -- either through the functional response or as intraspecific competition – dampens the expression of instability (Jensen & Ginzburg 2005). Alternatively, competitive modification of food web components (mean-driven ecosystem instability) can cause ecosystem malfunction through the addition of an inedible, or less edible, resource (Tilman 1982; Abrams and Walters 1996; Vos et al. 2004). Here, instead of boom and bust cycles in abundance, the inedible resource (e.g., primary producers laden with secondary chemical compounds) takes up the excess nutrients and flourishes. The flourishing inedible prey ultimately outcompetes the edible resource driving it to extinction (termed a transcritical bifurcation in dynamical systems theory; Grover 1995). Note, the result of inedible takeover is a specific instance of competition theory (Tilman 1982; Chase & Leibold 2003). This alternative route to POE leads to a massive loss in stability expressed as the complete replacement of the original ecological equilibrium with a new equilibrium due to a phase transition to a new state (Scheffer et al. 2001; Carpenter 2005; see Box 2). Ecosystem theory predicts large detrital accumulations in the new steady state as high resource pools die off and ultimately lead to the accumulation of large detrital pools, something we see frequently in the empirical examples (e.g., build-up of dead plant material in enriched grasslands; Tilman and Wedin 1991)).

In the 1990s Gary Polis and others championed the notion that nutrient and resource subsidies across ecosystem boundaries alter the structure and function of local ecosystems (Polis & Strong 1996; Polis *et al*. 1997; Leroux & Loreau 2008). These ideas produced theory that showed, for example, that nutrient subsidies can lead to dynamic outcomes that are stabilizing in the sense that they reduce boom and bust dynamics; however, they also again can drive strong top-down suppression such that consumers can often reduce their resources to low densities or extinction (Huxel & McCann 1998; Takimoto *et al*. 2002). This is not the destructive oscillations predicted by Rosenzweig’s POE, but rather another nutrient-driven loss of an equilibrium (i.e. mean resource densities suppressed to extinction).

The problem of nutrient run-off has also inspired models that predict alternate states in lakes and coastal ecosystems (Scheffer et al. 2001; Carpenter 2005). Here, though, the loss of an equilibrium is due to an abrupt shift to a new alternate ecosystem state through a saddle-node bifurcation. More recently, this nutrient theory has progressed to analyzing meta-ecosystems whereby the coupling among ecosystems creates feedbacks that can propagate instabilities. This cross-ecosystem research has added to DeAngelis’ seminal contributions that showed ecosystem models readily produced instability and POE (DeAngelis 1992). Intriguingly, one of the findings from meta-ecosystem research is that the paradox of enrichment, or strong consumer-resource oscillations, can be driven by nutrients coming from a neighboring ecosystem – this is the first theoretical suggestion that nutrients can spatially propagate instability across ecosystems (Gounand *et al*. 2014).

## A Synthetic Meta-Ecosystem Model: Nutrients and Consumers Connected in Space

In order to integrate the theory discussed above, we use a spatial ecosystem model (Fig. 2A-C). The base of this model is a fundamental ecosystem module that includes a detrital pathway and a consumer-resource interaction (DeAngelis 1992). This model is consistent with the clearest experimental evidence of the POE (Fussmann *et al*. 2000). The ecosystem module serves two main purposes: i) it allows us to follow the fate of nutrients locally and regionally; and ii) it allows us to monitor the local and regional stability properties of one of the major building blocks of food webs (i.e., consumer-resource interactions). Importantly, the consumer-resource sub-system of this module will allow us to synthetically bridge our model results to the historical theory discussed above (McCann 2011; Murdoch *et al*. 2013). While we employ a simple base module, we see it as a significant starting point for interpreting whole food web effects on the landscape. Qualitative general results from consumer-resource theory have been recently shown to scale coherently to whole food webs (Gellner and McCann 2016). We also extend our spatial model to include less edible prey, to show how food webs can alter the form of instability instigated by increased nutrients (Fig. 2C).

**Fig. 2.**
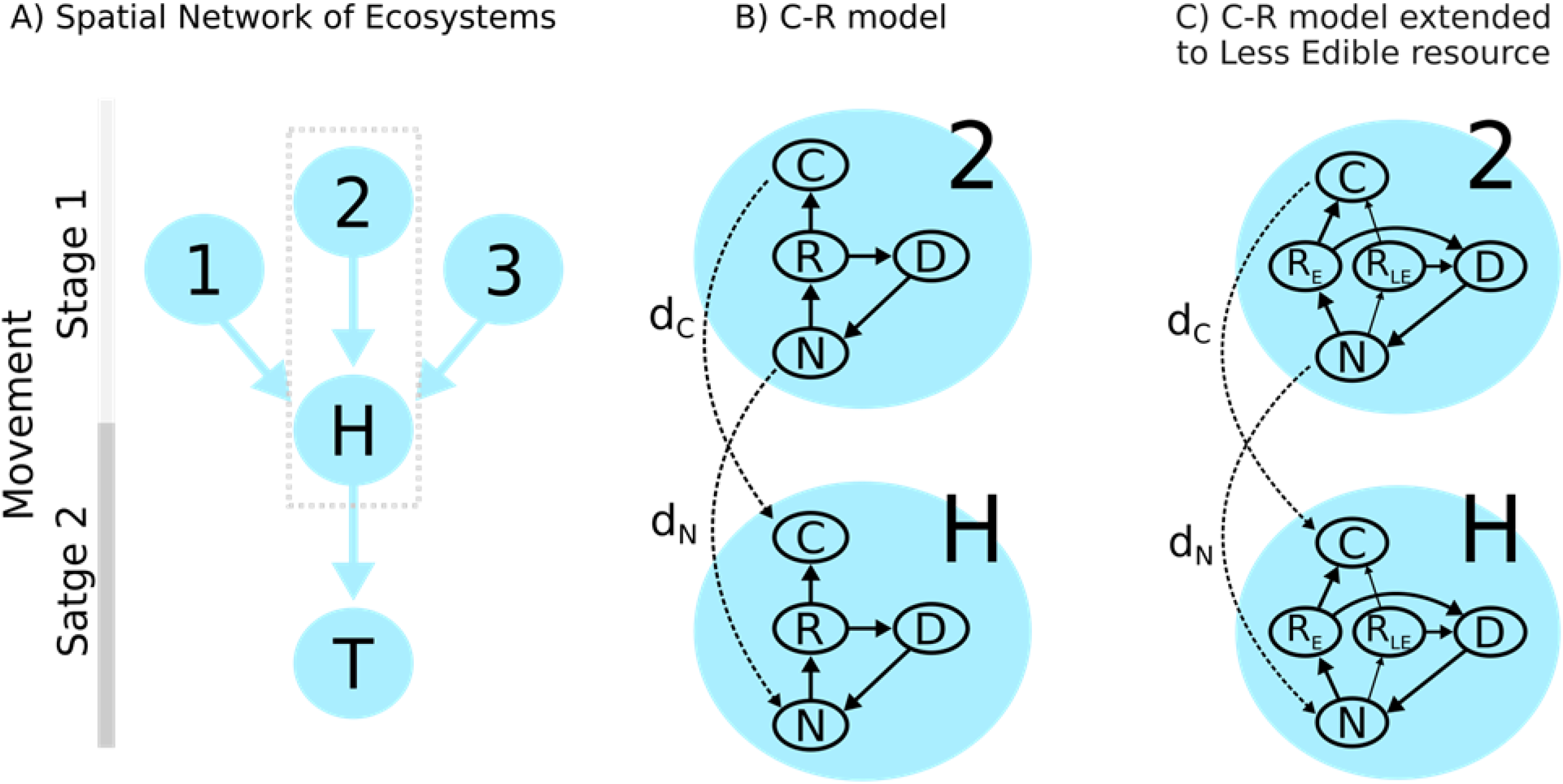
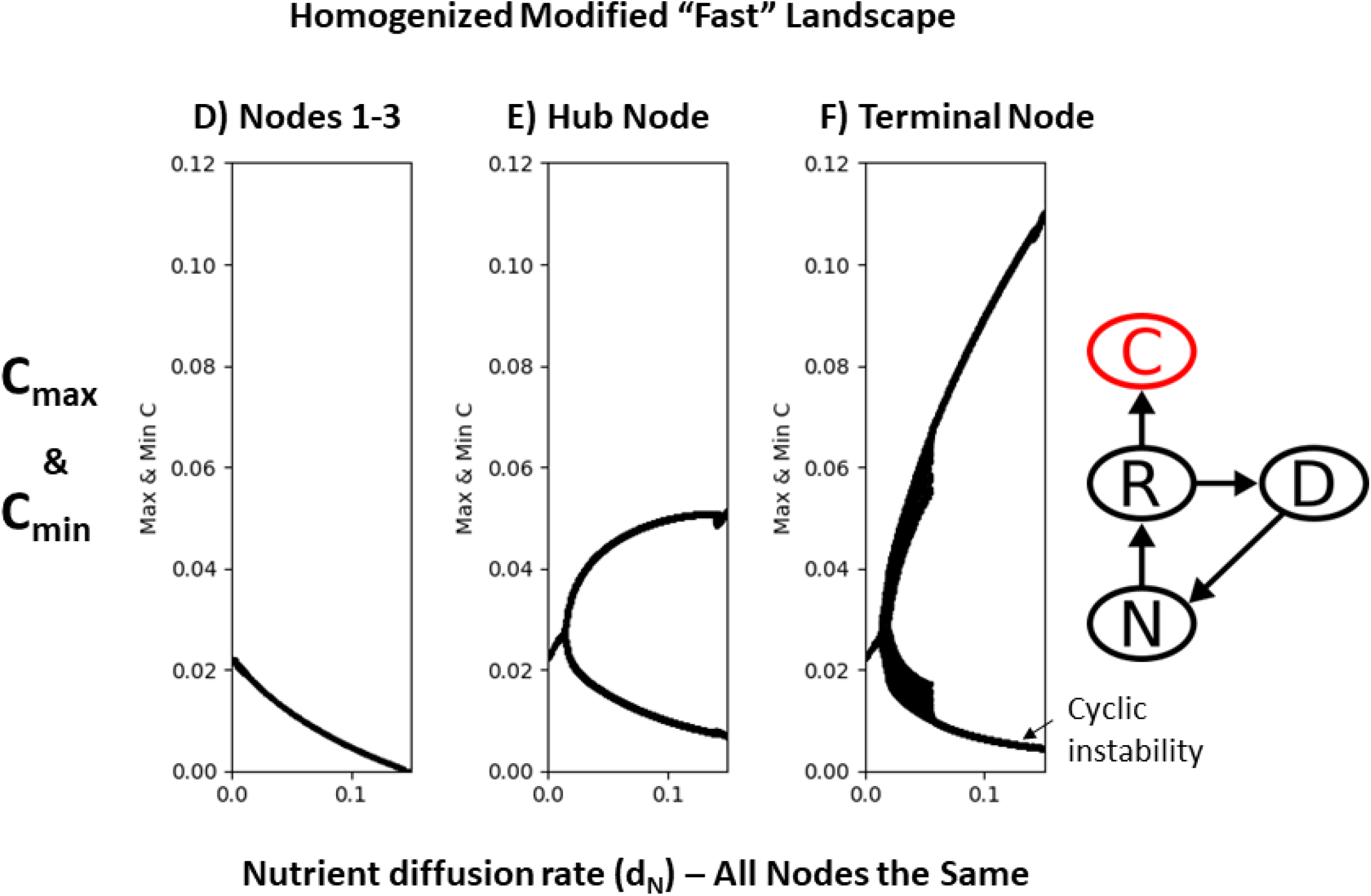
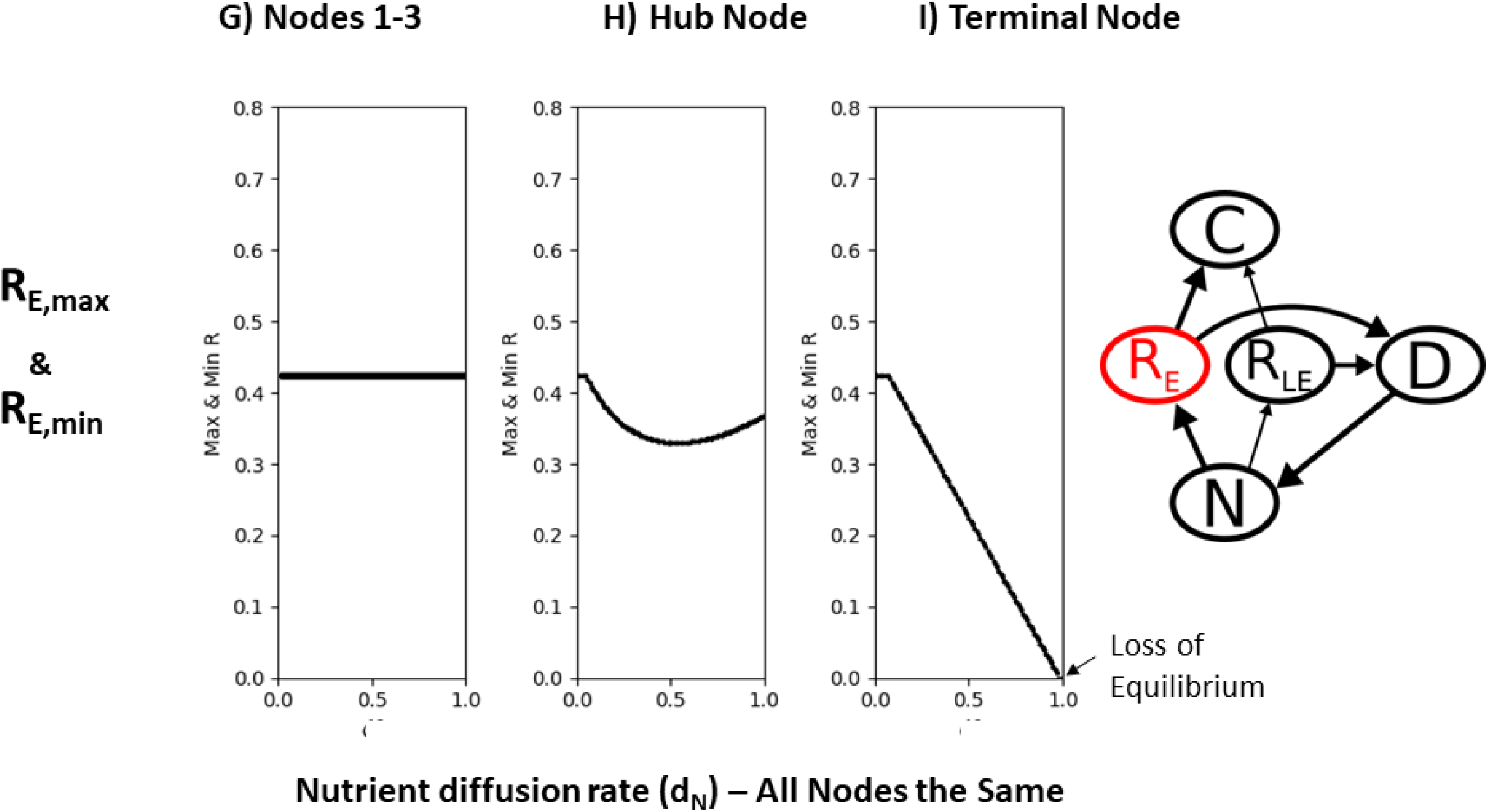
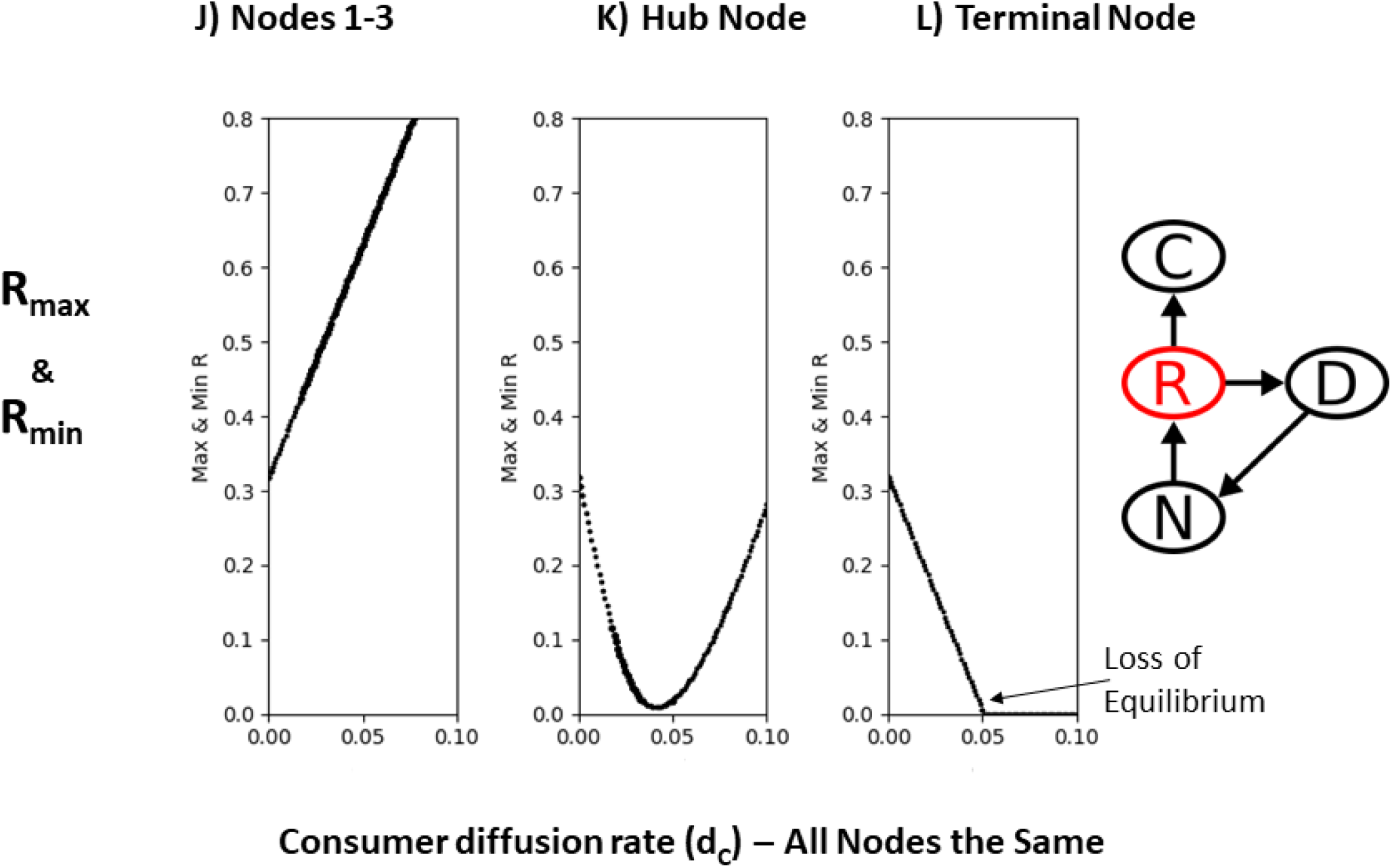
Simple dendritic or directionally connected model with multi-stage movement and an ecosystem model in each node. All nodes are considered “fast” and so all have high and equivalent nutrient loading, *I_N_*, and all nodes are changed with changes in nutrient diffusion, *d_N_*. A) The spatial arrangement with initial nodes (1, 2, and 3), an intermediate hub node (*H*) and a terminal node (*T);* B) a standard ecosystem module, and; C) the same ecosystem model extended to consider edible resources (*R_E_*) and less edible resources (**R_LE_**). Nutrient dominated movement results from model: D-F) Local max and min solution of the consumer, *C*, over a range of nutrient diffusion rates (*d_N_*) for nodes 1, 2 and 3 (all rates for initial nodes 1-3 are same in the homogenized case), the hub node and the terminal node for the standard ecosystem model. Results show oscillatory instability in terminal node (POE) relative to nodes 1-3. G-I) Local max and min solution of edible resource, *R_E_*, over a range of nutrient diffusion rates for node 1, 2 and 3, the hub node and the terminal node for the standard ecosystem model with both edible and less edible resources (all nodes *d_N_* are the same as all are fast nodes). This latter (G-I) result differs from the POE cyclic instability in that it shows that the takeover by inedible resource coupled to the generalist consumer ultimately pushes *R_E_* through a transcritical bifurcation (see Box. 1 for bifurcation definitions). This loss in stability is of a different flavor than the classic POE but nonetheless is a loss that fundamentally alters the ecosystem due to nutrient increases and nutrient transport. Consumer-dominated movement results from model: J-L) Local max and min solution of resource, *R*, over a range of consumer diffusion rates (*d_C_*) for nodes 1-3, hub node and the terminal node for the standard ecosystem model (again, all nodes experience *d_C_* as all are considered fast nodes). Note, *R* is suppressed to extinction here through a transcritical bifurcation and is a loss of stability due to the complete loss of an interior equilibrium.

To understand the role of spatial networks we assume a simple spatial unidirectional dendritic network (Fig. 2A) and a terminal node. The spatial network can be considered to have two stages of movement allowing us later to relate this theory to empirical results, which are often defined by multiple stages of movement and connectivity (Fig. 2A). As an example, the spatial model can mimic a simple dendritic network linking agricultural run-off from streams to rivers (stage 1 movement) and rivers to a terminal larger waterbody (stage 2 movement). Similarly, one can envision the spatial model as a network linking terrestrial nutrient flows to the ocean (stage 1 movement) and a second stage whereby ocean currents move the nutrients at intercontinental scales (stage 2 movement).

For simplicity, we assume that each spatial node has two potential parameter configurations: an unmodified pristine state with lower nutrient loading (*I_N_*) and reduced flows between spatial nodes (*d_N_* or *d_C_*) or a modified state with high nutrient loading (*I_N_*), and high flow rates (*d_N_* or *d_C_*) between neighboring nodes. When each node in the entire grid is in the modified state we are examining the impact of a homogenized landscape. In order to study stability, we employ local maxima and minima of population dynamics as it allows us to rapidly assess changes in population variation (i.e., consistent with classical POE) and it also simultaneously allows us to visualize structural changes in equilibria. This equilibrium-tracking allows us to monitor where we have instability through the complete loss of an interior equilibrium. The local maxima and minima thus allow us to see where enrichment in space drives cyclic instability (high variance with dangerously low minima) as well as changes in mean-driven ecosystem instability (Gellner *et al*. 2016).

We look at how increasing rates of movement through the nutrient-subsidized nodes impact the dynamics of the ecosystem module in space. We do this by increasing movement rates for nutrients (*d_N_*) or consumers (*d_C_*) and looking at the spatial structure of attractor solutions, as well as stability metrics for the deterministic system. Finally, we conduct these experiments under two movement scenarios: i) a **nutrient movement scenario** where nutrients dominate connectivity on the landscape (via *d_N_*), and; ii) a **consumer movement scenario** where consumers dominate connectivity on the landscape (via *d_C_*). Note, in both movement scenarios we are always considering increased local nutrient inputs (high *I_N_*) in the modified nodes on the landscape, so our model experiments always link nutrients to spatial stability.

## Homogenized Landscape, Rapid Nutrient Movement and Distant Instabilities

We first consider the case of a homogenized spatial model (all nodes are equivalent with high nutrient loading, *I_N_*) and we look at how altering the spatial connectivity of this landscape (via increasing diffusion rates, *d_N_*, of the nutrients) influences model solutions in space (model 1 in methods). Our first result is that under a homogenized modified landscape with nutrient movement dominating the connections between ecosystems, the terminal node in a directed spatial network eventually garners the most nutrients (Fig. 2). In this simple unidirectional flow model nutrients tend to build up in the terminal cell when flow rates between nodes grow (when at equilibrium this produces a surge of nutrients (i.e., equilibrium values of N that increase to maximum values in terminal node) in the dendritic network). In essence, the dendritic network under high connectivity yields a strong **spatial inflation of nutrients**. This inflated flux of nutrients across the landscape, consistent with the original POE, drives the greatest instability ultimately in the terminal node (Fig. 2D-F) as seen expressed with the increasing local max and local min for a given diffusion rate, *d_N_* (compare Fig. 2D, or node 1, versus Fig. 2F, or terminal node).

While an interesting result in and of itself, food webs are clearly not as simple as the ecosystem module above. One common natural component of many aquatic ecosystems is the presence of resources (e.g. algae) that are relatively inedible (model 2 in methods). This has the potential to change the stability result above or at least change the expression of instability. Fig. 2 shows the results for this model for an initial node (Fig. 2G), the hub node (Fig. 2H) and the terminal node (Fig. 2I) but this time follows edible resource densities since these densities can be altered strongly by the presence of a less edible resource competitor for nutrients. Again, there is a standing wave of nutrients (not shown here) but this time the edible resource is ultimately pushed to local extinction directly (i.e., it hits zero densities in a transcritical bifurcation; Fig. 2I) -- this is the second form of instability and different than the POE version above (Box 2). In the POE the dynamics drive wild oscillations with average consumer, *C*, densities increasing and average resource, *R*, densities staying steady. The solution in the POE case never actually reaches 0 but approaches it – in the real world these would very likely lead to local extinction of *C*, or both *R* and *C*. Instead, in this latter case, the mean density of edible resource (*R_E_*) is pushed lower until the equilibrium solution intersects with the *R_E_ = 0* axis. Effectively, less edible resources stabilize cyclic instability but give way to mean-driven ecosystem instability (i.e., high biomass of less edible resource). The responses of the other state variables are not shown but both less edible resources, *R_LE_*, and detritus, *D*, grow to high densities in the terminal node. High detrital biomass, *D*, correlates to high bacterial loads and increased oxygen consumption potentially threatening critical ecosystem function.

Our second result, then, is that nutrient-driven instabilities (i.e., loss in persistence of a species) in a homogenized landscape can occur at a distance through different bifurcations, here seen as the complete loss of an interior attractor. No matter which path leads to instability (e.g., Hopf versus transcritical), they have similar consequences for ecological systems -- **high local nutrient inputs shunted rapidly across the landscape drive instability and ecosystem imbalance in a distant terminal node**. Further, all cases above suggest that the increase of nutrients and their speed of transport over time lead to a phase transition to a new ecosystem state (at the onset of the nutrient-driven bifurcations). Note, our simple base ecosystem models generically produce transcritical outcomes but alterations to these models with positive feedbacks (e.g., discussed cogently by Scheffer et al. (2001)) would change these results to abrupt transitions to alternate states. Nonetheless, the general result holds – nutrients propagated rapidly drive phase shifts to a different equilibrium (e.g., a detritus-dominated equilibrium state) at great distances. These instabilities caused by rapid spatial movements of nutrients resonate with non-spatial ecological network results in that high flux through a food web, or strong interaction strengths, generally drive instabilities (see Box 3) suggesting that high flux scenarios in spatial or ecological networks are instability generators. Research on ecological networks has found that weak interactions can prevent these potent instabilities (structural energy deflection; see Box 3) and so this immediately suggests that a similar spatial result may hold.

### Box. 3 Non-spatial Muting of Enriched Food Web Networks: Structured Energy Deflection

It is informative to look at results from ecological networks to understand the outcome for stability in directional spatial networks. We briefly discuss theory that argues how food web structure can mute the instabilities discussed in Box 2. Specifically, the notion that weak interactions can stabilize both modular and whole food webs relies, to some extent, on the notion that certain common food web structures can re-route energy away from a strong interaction and in doing so stabilize a food web (where stability here can mean reduce CV either through reducing variance (*SD*) or reduce CV by increasing the mean (*μ*)). As an example (Fig. B.3.A), McCann et al. (1998) performed simple experiments where adding a weak interaction strength to a key interaction (e.g., reducing attack rate) on a wildly oscillatory food web module rapidly reduced the size of the oscillation, even potentially fully stabilizing the system (e.g., becomes stable equilibrium; CV=0). Here, we introduce a relatively inedible competitive consumer 2 into a wildly oscillatory P-C-R system and note it deflects energy away from a strongly interacting food chain. In doing so, it reduces the CV and stabilizes the dynamics. We will refer to this form of ecological network stabilization as **structural energy deflection** and note while it works in non-spatial ecological networks there is clearly room for a spatial analog as theory suggests high nutrient and energy flux in space ought to generally drive instability (we show that indeed it does).

This deflection of energy also can similarly impede the loss of an equilibrium (see Box. 2). As an example, we have a subsidy model driving strong top-down impacts on an intermediate consumer (Fig. B.3.B). This top down suppression with enough subsidy can remove the intermediate C1 (a form of mean-driven instability associated with a loss in equilibrium). If we introduce to this same subsidy a weak food chain (Fig. B.3.B), the subsidy is deflected away from top predator 1 (some portion going to the weak predator 2), removing the pressure on the intermediate consumer 1 and impeding the loss of stability (consumer 1 is relieved of top-down suppression). Again, as above, weak energy deflection away from the strong interaction removes the instability. Note, while this structural deflection works in non-spatial ecological networks **we conjecture that this same general form of material or energy deflection can play a potent role in stabilizing spatially connected food web networks but rather than deflecting energy into weaker species or food web pathways (as above) nutrient and material flow on the landscape is deflected in space away from strong nutrient pathways and thus inhibiting high nutrient aggregations in space**.

**Fig. B.3.**
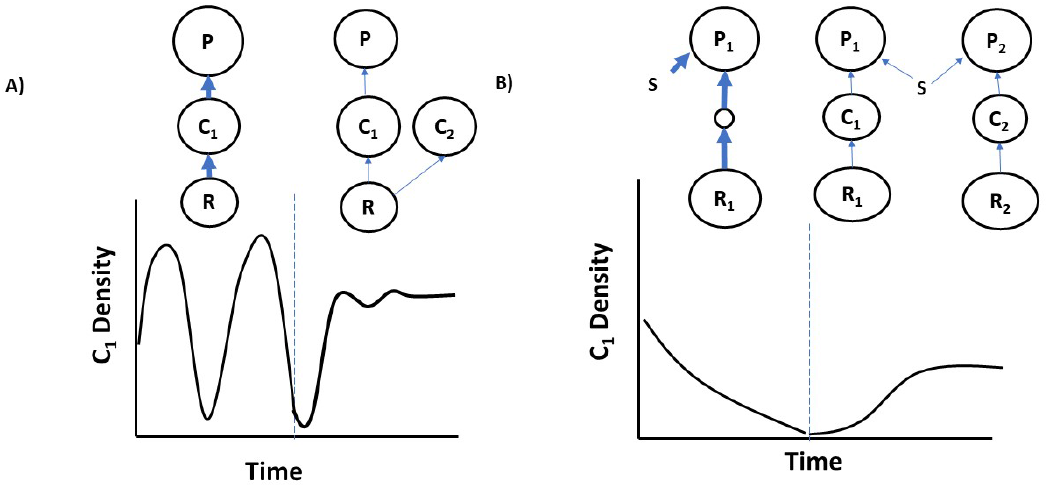
Ecological network, or food web, structures that can mute the two forms of instability. A) Weak or inedible consumer mutes oscillatory P-C-R, and; B) Weak food chain introduced and it deflects subsidy away from top predator in the strong food chain, and in doing so releases consumer suppression muting the loss of instability due to a complete loss of an equilibrium.

Next we consider how altering the spatial connectivity of this landscape (via increasing movement rates, *d_C_*, of the consumer) influences the model solutions across space (model 1 with consumer diffusion only) using the homogenized spatial model (all nodes are equivalent with high nutrient loading, *I_N_*). Under a homogenized landscape with consumer-movement dominating the connections between ecosystems, the terminal node in a directed spatial network garners the most consumers. What we see here then, akin to our first experiment, is the **spatial inflation of consumers**. Fig 2J-L show the dynamic changes in local max and min of the resource in node 1 (Fig. 2J), the hub node (Fig. 2K,) and the terminal node (Fig. 2L) to increasing consumer movement and connectivity. The terminal node inflation of *C* ultimately suppresses *R* (Fig. 2L) and drives the loss in stability through the complete loss of the interior equilibrium. This is similar in spirit to the inedible case above. For low *d_C_*, the hub node first accumulates consumers from the dendritic connections and suppresses *R_E_* before *d_C_* is fast enough to push consumers to terminal node. Indeed, although we have only looked at a simple set of models, the loss of stability due to high nutrients in the terminal node ought to be a very general response. It is worth commenting here on a relatively common empirical case (discussed below) whereby each stage of movement may be dominated by different facets of movement. Specifically, the first stage may be dominated by nutrient movement and the second by consumer movement. While the specifics of the model assumptions change, these results still lead to less stable terminal nodes that are some combination of the above two endpoint movement cases.

## Slow Nodes and Spatial Heterogeneity as a Potent Stabilizing Force

We end our development of the theory by considering the case of a heterogeneous spatial model (model 1) – we include both modified nodes with high nutrient loading (*I_N_*) and an unmodified, slow hub node with lower loading and diffusion rate (*d_N_*) into the terminal node, and we assess how altering the spatial connectivity of this landscape via increasing diffusion rates, *d_N_*, of nutrients from nodes 1-3 impact the model solutions in space. Here, we find that heterogeneity in the rate of diffusion stabilizes the terminal node (Fig. 3A-C) relative to the purely modified homogeneous case (Fig. 2D-F). More specifically, heterogeneity can alter where instability is expressed in the network (Fig. 3). Fig. 3B shows that nutrient inflation occurs in the now slow hub node, resulting in local instability, which then reduces the flux of nutrients to the terminal node, leaving it relatively stable (Fig. 3C).

**Fig. 3.**
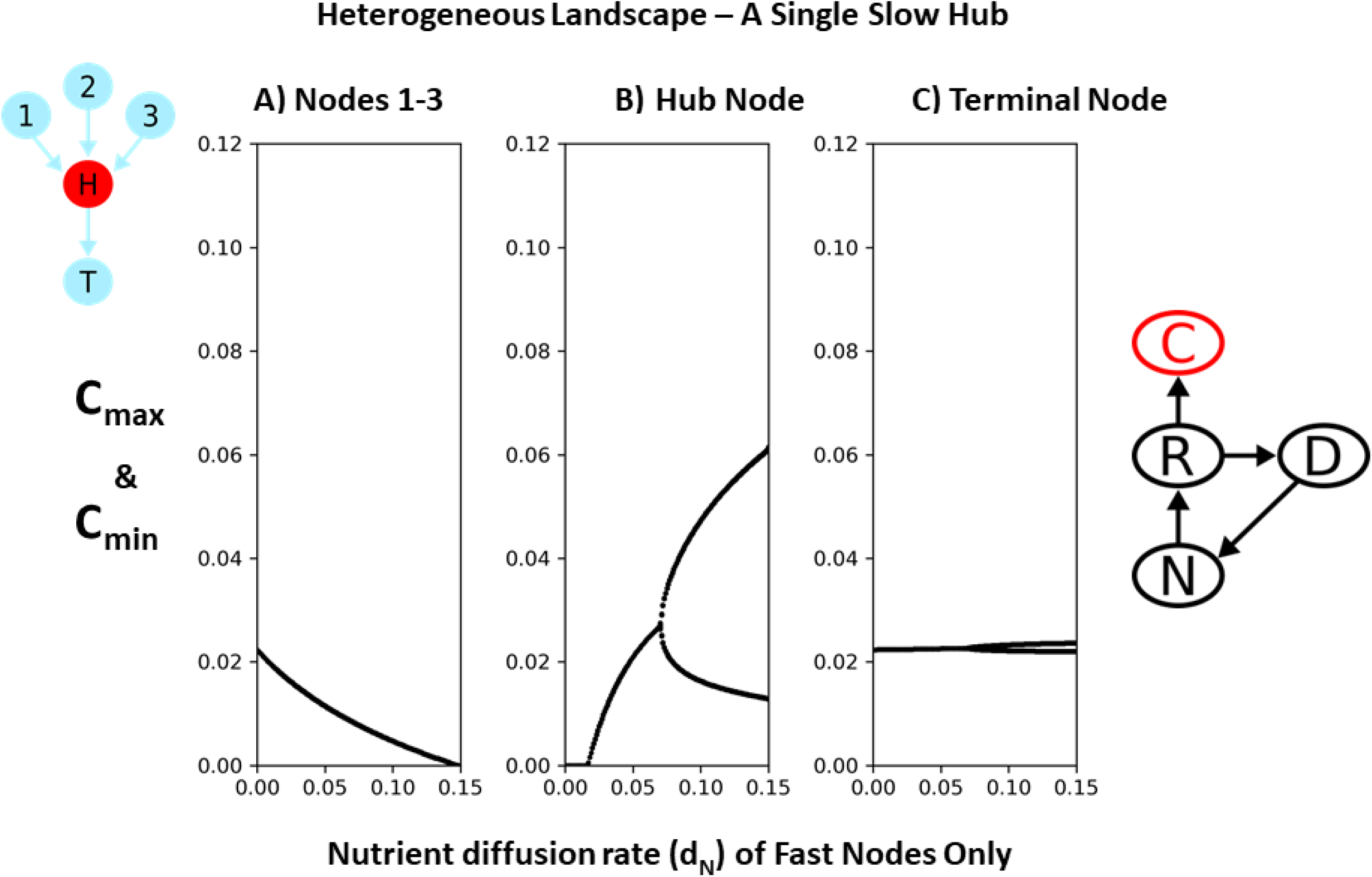
Example of how landscape heterogeneity that includes a node that slows flow rates or weakens landscape connectivity ultimately drives greater stability at terminal node. Deterministic min/max solution for increasing nutrient diffusion rates (*d_N_*) in nodes 1-3 while maintaining low *d_N_* and *I_N_* in the slow hub node (B). The terminal node response is also shown (C). In a manner similar to weak interaction theory (Box 3), the heterogeneity limits nutrient movements and alters the distant expression of instability as it is most unstable in the slow hub node.

This result highlights that strong connectivity drives instability at great distances, but that altering where the nutrients can be assimilated on the landscape determines how effectively nutrients can be “bled” off and thus preventing their effects from being expressed at the terminal node. Clearly, increasing the number and location of slow nodes on the landscape could reduce the stability of any one node while increasing the stability of the terminal node, and therefore overall regional stability by lessening the distant impacts of nutrient accumulation. This result resonates with the suggestion above, that low spatial flux – akin to weak interactions in ecological networks – can mute spatial instabilities (Box 3). An interesting extension of this work would be to understand how the near fractal pattern of riverine networks might alter stability by buffering flow rates. This issue of buffering flow rates not only relates to curbing distant accumulation of nutrients but has well-established practical implications for global efforts to “slow the flow” of floodwaters (Milly *et al*. 2002; Poff 2002). Those efforts seek to combat accelerating flooding with climate change, with flooding compounded by long-established barriers and channelization networks that, as with nutrients, sought to shunt water downstream as quickly as possible. Our work reinforces how these water management efforts, from field tiling to straightening rivers and modifying wetlands, solve local-scale concerns but are increasingly creating unanticipated regional and global scale impacts.

## Empirical Examples of Nutrient-Driven Ecosystem Imbalance

Empirical data show nutrients are consistently involved in long-distance ecological instability. Although some researchers have argued against the existence of empirical examples of cyclic dynamics from nutrient enrichment (Jensen & Ginzburg 2005), it is clear that nutrients have impacted ecosystem stability (Diaz & Rosenberg 2008; Smetacek & Zingone 2013; Hautier *et al*. 2014). Indeed, the most obvious and pervasive empirical example is the rising frequency of runaway harmful algal blooms (Diaz & Rosenberg 2008) (Fig. 4A). These events are often found in large lakes, seas and oceans (e.g., Lake Erie, Baltic Sea) – effectively terminal nodes of a spatial network (Diaz & Rosenberg 2008). These dominant inedible algae drive the mean densities of once prolific species (e.g., edible algae) to functional extinction, simultaneously diverting primary production from the rest of the normal food web and completely altering ecosystem structure. This structural or mean-driven instability appears as a competition-driven process whereby edible algae are reduced in density over time and replace by inedible algae due to increased levels of nutrient loading and enhanced nutrient coupling on the landscape. In its most extreme form, this mean-driven ecosystem destabilization can lead to dead zones, where top consumers in the aquatic food web have gone locally extinct. This empirical result resonates with the abrupt loss of an equilibrium or equivalently a phase transition to a new diminished ecosystem equilibrium state discussed in the spatial nutrient-stability theory above (particularly model 2).

**Fig. 4.**
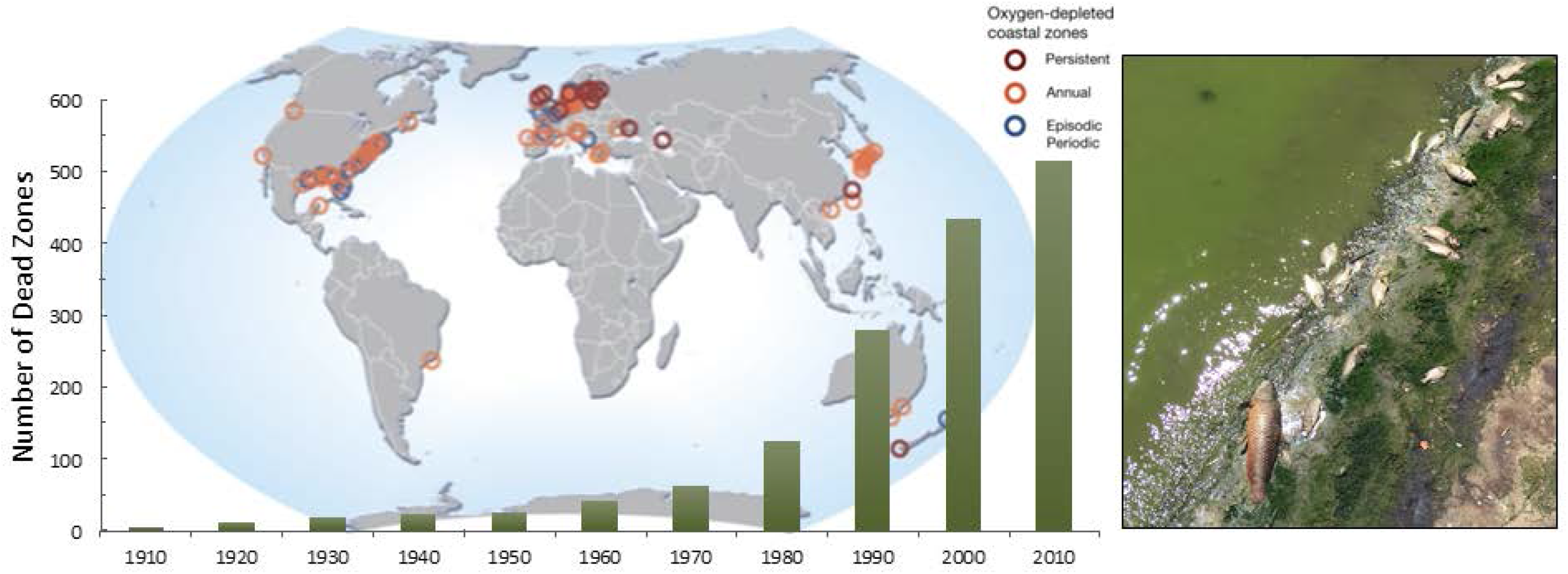
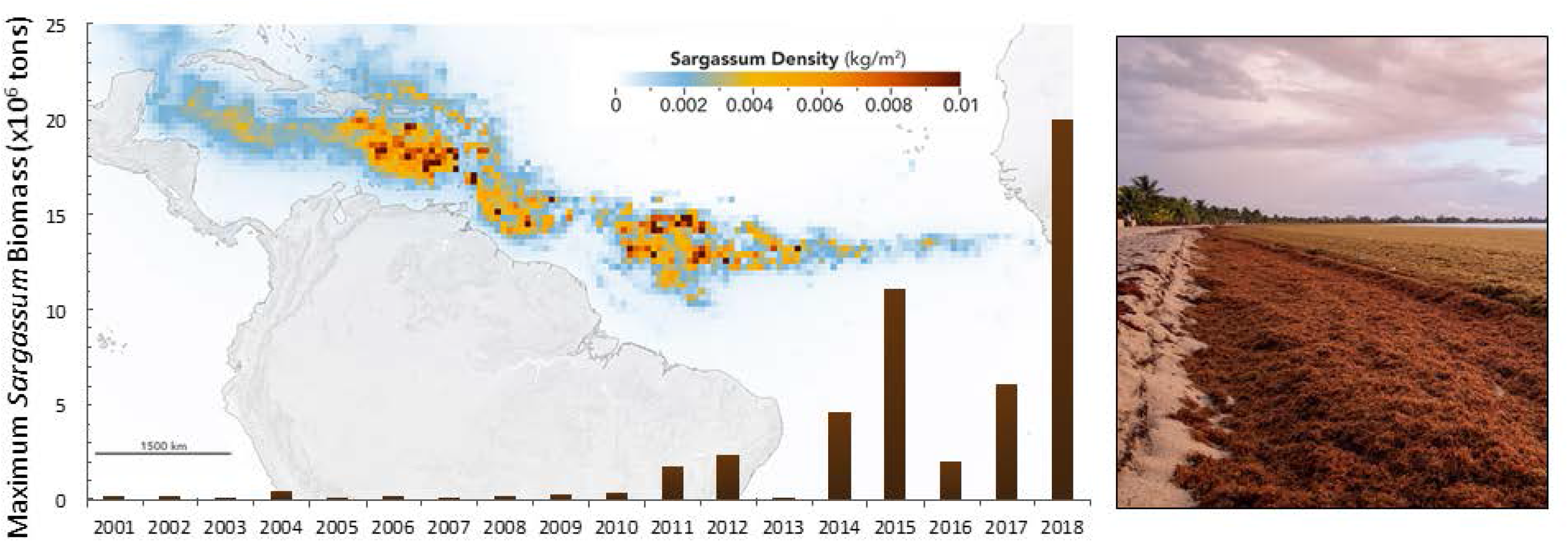
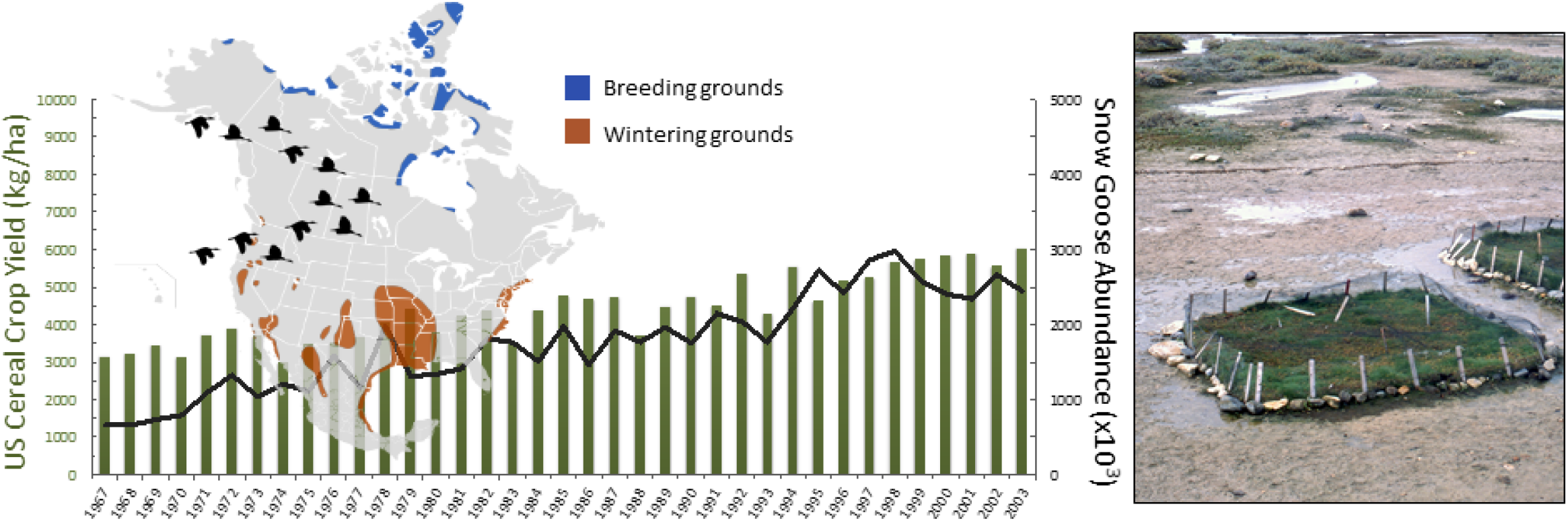
Some examples of nutrient-driven instabilities at a distance. A) Dead zones of nutrient and climate fueled runaway plankton followed by deep water bacterial growth that removes oxygen, both in large freshwater lakes and oceans, have been noted to increase in intensity and frequency globally; B) Similarly, nutrient and climate fueled macroalgal blooms have also been increasing. In this example, *Sargassum* densities have been magnified by deforestation and nutrient loss off the coast of Brazil resulting in increasing intensity and frequency of blooms that have been swept thousands of kms around the ocean landing on terrestrial coastal zones; C) A terrestrial mobile consumer, snow geese, has grown in step with the use of increased nitrogen on cereal crops in the southern USA to migrate thousands of kms North where it denudes whole marshland ecosystems. A) Map from: Hugo Ahlenius, UNEP/GRID-Arendal Time series data from: Convention on Biological Diversity Secretariat 2010 Photo credit: Kirsten Macintyre, California Department of Fish and Wildlife B) Map from: NASA/Earth Observatory. Data provided by Mengqiu Wang and Chuanmin Hu, USF College of Marine Science Time series data adapted from: Wang et al. 2019 C) Map from: Cephas, CC BY-SA 3.0 https://commons.wikimedia.org/w/index.php?curid=5980177 Snow geese data from: Jeffries et al. 2004 US Cereal crop yield from: Food and Agriculture Organization Photo credit: Peter Prokosch www.grida.no/resources/4428

Harmful algal blooms and aquatic dead zones have been argued to be driven by nutrient run-off from agriculture and urban development, coupled to increased temperatures that further accelerate algal growth rates, suggesting that the rising tide of imbalance may be created synergistically by increasing nutrients and climate change (Diaz & Rosenberg 2008; Huisman *et al*. 2018; Ho *et al*. 2019). Yet climate change may do more than increase growth rates. It has been suggested that, on top of the spatial amplification of nutrients in dendritic landscapes, climate change has also been associated with increased temporal variation in meteorological events that can temporally amplify nutrient run-off (e.g., during a large rainfall event) in time and space (Michalak *et al*. 2013). Scientists have argued that in the Lake Erie agricultural basin, the extreme spring rainfalls or snowmelt can rapidly carry off fall or winter-applied fertilizer, creating highly concentrated nutrient pulses that ignite blooms to record levels (Michalak *et al*. 2013; Motew *et al*. 2018). Here, spatial and temporal nutrient inflation combine to intensify the nutrient-driven ecosystem imbalance at a distance.

Interestingly, little work has been done in agriculturally dominated landscapes to look at where in a spatial network we experience blooms or ecosystem imbalance. The empirical results above suggest that wherever we have a slowing down of nutrient transport, which in turn increases nutrient deposition, we may expect increased biotic nutrient assimilation accompanied by nutrient-driven imbalance. Recent work looking for mini-blooms in an urban streams network have found outbreaks of harmful algal blooms where storm events have scoured the streams to set up slow moving stream pools (Blaszczak *et al*. 2019). Here, the combination of fast, high nutrient transitions into these slow pools, even small ones, are a recipe for resource blooms as rapid nutrient movement reduces efficient biotic assimilation of excess nutrients until it hits the slow pool where nutrients are assimilated, and instability is expressed. Additionally, there can be significant impacts on coastal ecosystems like seagrasses or coral reefs. For example, a 30-year long-term data set on nutrients and climate found that while temperature stress causes coral reef bleaching, the impacts are exacerbated by an interaction with nutrient loading (Lapointe *et al*. 2019). Sewage, fertilizers and top soil from a large region of Florida inflate nitrogen levels entering the Keys hundreds of kms away, which then alter the coastal N:P stoichiometry ultimately starving corals of phosphorus, and in turn reducing their temperature threshold for bleaching (Lapointe *et al*. 2019). Note, this case acts as an unintentional ecosystem entanglement experiment. Algae blooms correlate perfectly in time with the sudden speeding up of water movement by the Army Corps of Engineers – an action to reduce hyper-salinity in Florida Bay – suggesting a strong coherence with the ecosystem entanglement ideas presented here. This result suggests that the cascading impacts of local actions are likely empirically underestimated as they potentially move across many connected ecosystems.

On a similar empirical note, *Sargassum* – a pelagic macroalga – has risen in density over the last eight years and appeared almost magically stretching across the tropical Atlantic Ocean (Wang *et al*. 2019). Over this time period, the macroalgae have been accumulating in vast quantities on beaches throughout the Caribbean and Central America to the consternation of coastal tourism. The origin and reason for this large-scale spatial bloom was unclear until recent work using global satellite imagery revealed yet another ecosystem imbalance at a distance (Wang *et al*. 2019). Land cleared in Brazil, and subsidized with nutrients for agricultural development, increased the landscape connectivity and, in turn, water flows with the nearby ocean becoming the recipient of excess nutrients. Further, this increased connectivity is exacerbated by climate change producing extreme rainfall events and flooding of the Amazon basin (Wang *et al*. 2019). In turn, the pelagic macroalgae harness these human-driven inputs and bloom off the coast in the plume of the terrestrial-derived nutrient runoff (Fig. 4B). As modeled in the theory section above, there are arguably two major stages to this ecosystem connectivity with the first stage driving nutrients from a denuded landscape into the pelagic waters off the coast where *Sargassum* multiplies (Fig. 4B). The bloom (i.e., the resource) is then connected in the second stage of movement by large scale oceanic currents that sweep the resource at massive spatial scale (thousands of kms) to distance coastal regions (Wang et al. 2019; Fig. 4B). This is a poignant case of local actions on the land being coupled to nearby coastal ocean ecosystems, with oceanic conveyor belts carrying the impacts over the Atlantic basin, Caribbean Sea, and Gulf of Mexico, causing ecosystem imbalance at a distance. This macroalgal ecosystem instability is not isolated. Other instances include the macroalga *Ulva*, which has experienced runaway growth from nutrient run-off and climate in China (Liu *et al*. 2016) and effluents in Brittany (Charlier *et al*. 2006). Collectively scientists have called these macroalgal blooms green and golden tides and consistent with plankton blooms noted their increasing frequency (Smetacek & Zingone 2013).

Terrestrial examples also exist that show these distant impacts. Snow geese foraging on cereal crops in the southern USA have attained extremely high densities (Jeffries et al. 2004; Fig. 4C), corresponding in sync with the agricultural use of industrially made nitrogen accelerating in the 1950s near the beginning of the “Green Revolution” (Jefferies et al. 2004). Subsidized to high densities from these nutrient-fueled cereal crops, the snow geese migrate thousands of kms to the Hudson Bay lowlands during the summer. Here, they collectively overgraze marshlands leaving these once complex ecosystems as denuded hypersaline mud flats (Jefferies et al. 2004). This example emerges as another empirical case of the complete loss of an equilibrium, or mean-driven ecosystem instability (i.e., the marsh sedge resources are totally suppressed) but here the suppression occurs through spatial consumer inflation in the far north triggered by localized nutrient additions thousands of kms away to the south. Here, nutrient-driven instability is facilitated by the movement of inflated consumer densities, akin to our theoretical result above. In the case of snow geese, this result is exacerbated by positive feedbacks in the terminal node that make it even harder to potentially remediate these distant impacts of nutrients.

## Discussion

Physicists have noted “quantum entanglement” whereby two particles are paired instantly and remain intimately connected across vast distances. Here, using simple ecological theory, we show that ecological systems can become causally entangled over great distances (i.e., strongly connected), resulting in dramatic ecosystem imbalances. Although this entanglement is not instantaneous, it occurs rapidly, and the effects are often large in magnitude. With continued human modification of landscapes and waterways, this entanglement is becoming increasingly rapid and intensified. These non-local effects are frequently accompanied by great economic costs to society through the loss of critical ecosystem services (e.g., loss in water quality, secondary productivity, altered fisheries, tourism, degraded human health; Carpenter and Biggs 2009). The distant expression of local actions effectively couples cause and effect over space, and in the process makes for “wicked” environmental problems (Balint 2011) as they emerge from the cumulative effect of multiple isolated management decisions (e.g., fertilization, land clearing, river straightening) and across distinctly different political jurisdictions or regions (impacts hundreds to thousands kms away – e.g., Great Atlantic Sargassum Belt).

Our theoretical synthesis suggests that factors that increase directional connectivity (movement of nutrients or consumers) ultimately inflate the realized nutrient-driven instability in the terminal spatial node. As noted above, this spatial amplification of signals may be enhanced by climate variation (e.g., extreme rainfall) if it increases environmental variation that amplifies nutrient transport (Nelson *et al*. 2013). Arguments for this occurring in the magnification of the Lake Erie dead zone have already been made (Paerl & Scott 2010). The corollary to the nutrient inflation effect is that any spatial or temporal structure that reduces this hyper-connectivity can remove the distant instability (see Fig. 3 above) by bleeding off nutrients before they reach the terminal node. In a complex network, the diffuse assimilation of nutrients within the regional landscape can prevent distant nutrient inflation and costly ecosystem imbalance. We have kept our theory simple with the aim of motivating more detailed work on these directional movements that are major structural attributes of agriculture’s interaction with nature’s transport systems, such as streams and rivers (Fagan 2002; Grant *et al*. 2007; Peterson *et al*. 2013b; Moore *et al*. 2015; Terui *et al*. 2018). Ultimately, application of our results will require ecoinformatic approaches that combine detailed local and regional data with spatial modelling to inform how “slow nodes” can be placed effectively within a real network.

While we have focused here on the landscape effects of nutrient additions, clearly local nutrient-driven impacts (i.e., on the farm) can still have severe economic and environmental consequences (Carpenter et al. 1998). One might argue that distant effects stall management intervention because they are hidden far from view, or the impacts are observed but the origins unclear (e.g., it took almost a decade to determine the cause of the *Sargassum* bloom). Most fundamentally however, nutrient-derived instability, local or global, stems from dramatic inefficiencies in how global societies deal with nutrients whether farm applied, or from industry or urban sources (Carpenter et al. 1998; Huisman et al. 2018). For food production, these inefficiencies are part of a massive nutrient “retention shortfall where up to 20-30% of applied nutrients can miss their intended target (uptake by maturing crops) and flow off the farm, often because they are applied weeks or months before peak growth that, in turn, makes them susceptible to loss by extreme rainfall or spring melts. Efforts to reduce connectivity may ameliorate distant destabilization, with local destabilization possibly the lesser of two evils, but ultimately only reduced nutrient inputs from human-derived sources can truly control the problem (Zhang et al. 2015; Oita et al. 2016).

Here, we have concentrated on impacts of nutrient loading and its effects on the instability of ecosystems connected by dendritic networks. We focused on exploring the dynamical outcomes (i.e., cycles, loss of interior attractors) and the role nutrients and consumer movements play in mediating instability. As a result, we did not deal with the role of stabilizing portfolio effects whereby nutrients or populations that join different branches of spatial movement can average out variation (Tilman *et al*. 1998; Schindler *et al*. 2010; Thompson & Gonzalez 2016; Thompson *et al*. 2017). This averaging occurs strongly when different variabilities in different branches are asynchronous. Schindler identified a poignant example with sockeye salmon that return from different streams into a common ocean basin (Schindler *et al*. 2010). Clearly, future work on these dendritic networks needs to also consider how synchronizing forces across the network can disrupt stability. For example, regional spatial homogenization from agriculture likely increases the similarity of abiotic and biotic conditions across the network (e.g., all have high nutrients, all are channelized and fast, all have similar species) resulting in the loss of stability.

There is a growing number of examples of instabilities arising from the coupling of ecosystems across great distances (Diaz & Rosenberg 2008; Smetacek & Zingone 2013). Our framework articulates how landscape heterogeneity in nutrient loading and transport can mitigate these instabilities by allowing nutrient flows to be reduced and assimilated before reaching the ultimate end point. Our theoretical synthesis calls for a concerted research effort to understand how ecosystem entanglement occurs, and how it may be managed. The science we summarize here suggests that local actions have global consequences. N and P are both critical drivers of primary production, and support global agriculture and population growth yet ironically, they have also become major ecological pollutants. We have extended the POE from the local to the regional and global scales and suggested how the resulting ecosystem instabilities may be understood and mitigated. Specifically, we find that rapid nutrient transport leads to massive distant ecosystem instability, but these imbalances can be mitigated by the correct placement of slow nutrient transport nodes in the landscape network. Similarly, imbalances can be magnified by the incorrect placement of fast nodes. Given the social and economic costs of this ecosystem degradation, there is an opportunity to orient research and agricultural policy to manage the considerable environmental externalities arising from ecosystem entanglement. Collectively, this synthesis indicates that the paradox of nutrient enrichment is indeed real as Rosenzweig suggested nearly 50 years ago (Rosenzweig 1971) albeit this may be in more complicated ways as we move towards a more holistic landscape view of nutrient-stability theory.

## Acknowledgments

This project is funded by the University of Guelph’s Canada First Research Excellence Fund project “Food from Thought”.

## Author contributions

All authors contributed to the development of the idea through workshops and discussions. All authors contributed to the writing of the manuscript. KM, KC, GG and CB contributed to the development of the theory and coding. KM, AG and CB created the figures and boxes. AG is supported by the Liber Ero Chair in Biodiversity Conservation.

## Materials and Methods

### Models

The mathematical models were coupled ordinary differential equations that used simple ecosystem models as per DeAngelis (DeAngelis 1980). Spatial connections were done simply in a uni-directional way with the configuration given in Fig. 2A. Either nutrients moved passively (i.e., depart from node at a constant rate of their density via Fickian diffusion). Given this, the C-R ecosystem model (Fig. 2D-F, Fig. 2J-L, and Fig. 3) where node i is one of the **initial spatial nodes** (where *i={1,2,3}*) and is defined as (**MODEL 1**):

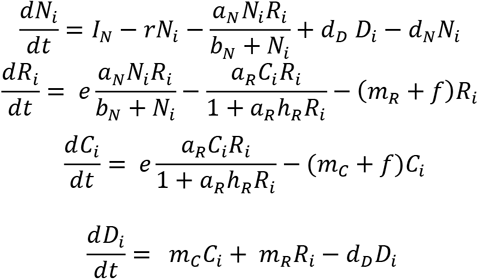

and **the intermediate node or the hub node (*H*)** is defined as:

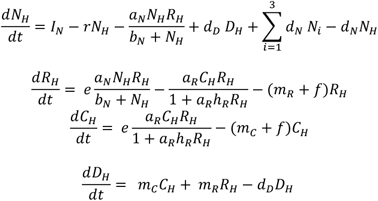

and the **terminal node (*T*)** is defined as:

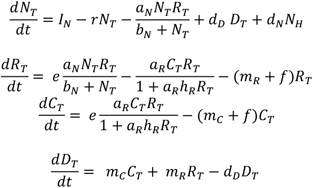

where *I_N_* is nutrient loading rate, *r* is nutrient loss rate due to leaching, *a_N_* is maximum nutrient uptake rate by *R*, *e* is conversion efficiency, *b_N_* is half saturation rate for nutrient uptake, *d_D_* is the rate of decomposition from detritus to nutrients, *a*_R_ is consumption rate of *C* on *R*, *h_R_* is handling rate of *C* on *R*, and *m_j_* is mortality rate of species or trophic level *j*, *f* is loss not recycled through the detrital pool, *d_N_* is the diffusion rate of nutrients between nodes.

The ecosystem model with edible and less edible resources includes a relatively inedible or less edible resource (Fig. 2G-I) Given this, the C-R ecosystem model (Fig. 2D-F, Fig. 3) where node *i* is the **initial spatial nodes (i={1,2,3})** and is defined as (**MODEL 2**):

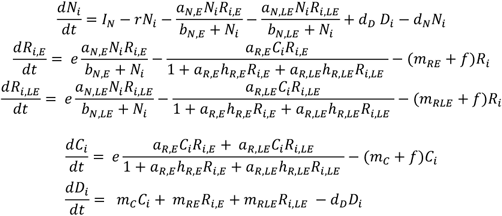

and **the intermediate node or the hub node (H)** is defined as:

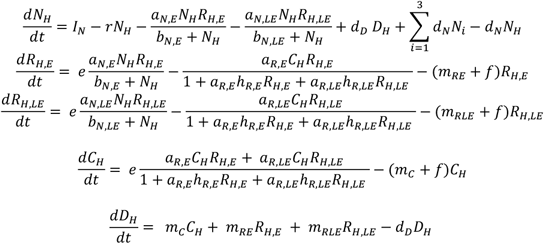

and the **terminal node (*T*)** is defined as:

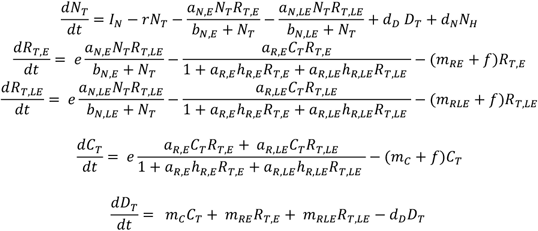

Where all parameters are defined above but now we have separate attack rates and handling times on a less edible resource (*R_LE_*) as well.

Finally, the ecosystem model with consumer movement (Fig. 2J-L) is the same as model (1) above but the C moves following passive Fickian diffusion (*d_C_*) not the resources.

### Methods

We ran all simulations of the above coupled ordinary differential equations for a transient of 5000-time steps before collecting local minima and maxima on the attractor for 5000-time steps for each value of the diffusion rate *d_N_* and then *d_C_*. Code was produced in the Julia programming language. These equations were often stiff and so we employed the KenCarp5 ODE solver.

### Parameters

Fig. 2. All models run for figure 2 assumed fast node parameters (i.e., high *IN*) and have the following parameters: Fig. 2G-I (model 2); Fig. 2J-L(model 1); Fig. 3 (model 1) All nodes assumed fast node parameters except the hub node which has slow node parameters:

**Fig. 2D-F (model 1)** – All nodes -- *IN=0.25, r=0.05, e=0.80, f=0.05, aN=0.60, bN=0.04, d=0.10, aR=0.20, hR=0.005, mR=0.10, mC=0.001*.

**Fig. 2G-I (model 2)** – All nodes --*IN=0.25, r=0.05, e=0.80, f=0.05, aN,E=0.40, bN,E=0.04, aN,LE=0.07, bN,E=0.047, d=0.10, aR,E=0.15, hR,E=0.005, aR,E=0.05, hR,LE=0.005, mR,E=0.10, mR,LE=0.11, mC=0.001*.

**Fig. 2J-L (model 1)** – All nodes -- *IN=0.25, r=0.05, e=0.80, f=0.05, aN=0.40, bN=0.04, d=0.10, aR=0.20, hR=0.005, mR=0.10, mC=0.001*.

**Fig. 3 (model 1)** – Hub Node -- *IN=0.09* all other parameters identical to Fig. 3.D-F; Fast Node – al parameters identical to Fig. 3D-F.

